# Necessity and recruitment of cue-specific neuronal ensembles within the basolateral amygdala during appetitive reversal learning

**DOI:** 10.1101/2022.03.08.483474

**Authors:** Sara E. Keefer, Gorica D. Petrovich

## Abstract

Through Pavlovian appetitive conditioning, environmental cues can become predictors of food availability. Over time, however, the food, and thus the value of the associated cues, can change based on environmental variations. This change in outcome necessitates updating of the value of the cue to appropriately alter behavioral responses to these cues. The basolateral amygdala (BLA) is critical in updating the outcomes of learned cues. However, it is unknown if the same BLA neuronal ensembles that are recruited in the initial associative memory are required when the new cue-outcome association is formed during reversal learning. The current study used the Daun02 inactivation method that enables selective targeting and disruption of activated neuronal ensembles in *Fos-lacZ* transgenic rats. Rats were implanted with bilateral cannulas that target the BLA and underwent appetitive discriminative conditioning in which rats had to discriminate between two auditory stimuli. One stimulus (CS+) co-terminated with food delivery, and the other stimulus was unrewarded (CS−; counterbalanced). Rats were then tested for CS+ or CS− memory retrieval and infused with either Daun02 or a vehicle solution into the BLA to inactivate either CS+ or CS− neuronal ensembles that were activated during that test. To assess if the same neuronal ensembles are necessary to update the value of the new association when the outcomes are changed, rats underwent reversal learning: the CS+ was no longer followed by food (reversal CS−, rCS−), and the CS− was now followed by food (reversal CS+; rCS+). The group that received Daun02 following CS+ session showed a decrease in conditioned responding and increased latency to the rCS− (previously CS+) during the first session of reversal learning, specifically during the first trial. This indicates that neuronal ensembles that are activated during the recall of the CS+ memory are the same neuronal ensembles needed for learning the new outcome of the same CS, now rCS−. Additionally, the group that received Daun02 following CS− session was slower to respond to the rCS+ (previously CS−) during reversal learning. This indicates that neuronal ensembles that are activated during the recall of the CS− memory are the same neuronal ensembles needed for learning the new outcome of the same CS. These results demonstrate that different neuronal ensembles within the BLA mediate memory recall of CS+ and CS− cues and reactivation of each cue-specific neuronal ensemble is necessary to update the value of that specific cue to respond appropriately during reversal learning. These results also indicate substantial plasticity within the BLA for behavioral flexibility as both groups eventually showed similar terminal levels of reversal learning.

**Highlights:**

- Chemogenetic inactivation of BLA neuronal ensembles activated by learned CS+ or CS−
- Examined if specific ensembles needed when cues’ values change in reversal learning
- CS+ ensemble ablation reduced responding to the same cue in early reversal learning
- CS− ensemble inactivation slowed learning of the new value of the cue

## 1. Introduction

Environmental cues can become strongly associated with food if they frequently occur together, and subsequent presentation of these learned cues can lead to food procurement and consumption without hunger (Birch et al., 1989; Holland and Petrovich, 2005; Petrovich, 2013; Petrovich and Gallagher, 2003; Saper et al., 2002; Weingarten, 1983). However, the outcomes, and thus the values, of associated cues are not always static and can change based on environmental variations. This change in the outcome of a learned cue requires updating the value of the cue to produce appropriate behavioral responses.

The basolateral nucleus of the amygdala (BLA) is a critical forebrain region necessary for associative conditioning and is an early processor of appetitive learning (Cole et al., 2013; Piette et al., 2012). The BLA is critically involved in appropriate behavioral responding when the values of learned appetitive cues are changed (Corbit and Balleine, 2005; Nomura et al., 2004; Schoenbaum et al., 1999; Tye et al., 2010; Fisher et al., 2020) or additional cues are incorporated to update the value of learned appetitive cues (Blundell et al., 2001; Everitt et al., 2003; Hatfield et al., 1996; Holland et al., 2002, 2001; Holland and Petrovich, 2005; Ishikawa et al., 2008; Petrovich, 2013; Setlow et al., 2002; Wassum and Izquierdo, 2015; Hoang and Sharpe, 2021). *In vivo* recording studies have shown that BLA neurons respond to appetitive cues, but then change their response profiles when cue outcomes are reversed (Schoenbaum et al., 1999) and when the reward is omitted (Tye et al., 2010). These studies suggest that the BLA neurons can alter their response to food predictive cues when the outcome changes. It is unknown, however, if the same or different BLA neuronal ensembles that are activated during the recall of learned cues are necessary for updating the values of these cues to form a new association when the outcomes change.

To this end, we used the Daun02 chemogenetic inactivation method (Cruz et al., 2013; Koya et al., 2009) to target BLA neuronal ensembles that are selectively activated by either the CS+ or CS− to determine if these specific neuronal ensembles are necessary to update the new value of the CSs during reversal learning. Specifically, *Fos-lacZ* transgenic rats underwent discriminative conditioning and were then infused with Daun02 or a vehicle solution into the BLA after presentation of either the CS+ or CS− to inactivate the responsive neuronal ensembles. Afterwards, rats underwent either one or fifteen sessions of reversal learning to observe how BLA neuronal ensemble inactivation affected conditioned responding to the initial memory recall of the CSs and during complete reversal learning, respectively. We hypothesized that separate BLA neuronal ensembles are activated during CS+ and CS− memory recall, and inactivating the neuronal ensembles that respond to a particular CS will only alter the memory of that CS and not the other CS. We also predicted that neuronal ensembles that are activated by a particular CS would be necessary to learn the new associations to the same CS when the outcome is changed during reversal learning.

## 2. Materials and Methods

Rats underwent surgery for implantation of bilateral cannulas aimed at the basolateral amygdala (BLA). After recovery, rats underwent ten sessions of discriminative conditioning followed by reversal learning as previously described (Cole et al., 2017; Keefer and Petrovich, 2020). Briefly, each session included 6 presentations of an auditory CS+ paired with the delivery of two food pellets (US) and 6 presentations of a separate, distinct auditory CS− presented alone. Following successful discrimination, rats underwent a brief induction session, which involved 6 presentations of either the CS+ or CS−, followed by infusion of Daun02 or vehicle ninety minutes after the beginning of the session. This induction session ensured selective activation of one CS neuronal ensembles without activating neurons recruited by the other CS or to the US. Next, rats underwent reversal learning where the outcomes of the CSs were reversed. Half of the rats were perfused 90min after the cessation of the first reversal session (R1) for histological verification of β-gal decrease in Daun02 infused rats (See Supplemental Materials). The other half received 15 reversal sessions to observe value updating after Daun02 inactivation. The primary measures of learning were the percentage of time rats expressed food cup behavior during the CSs and latency to approach the food cup during the CSs. Brain tissue was processed for double-label fluorescence immunohistochemistry for Fos and β-gal detection (see Supplemental Material and Methods for details). Notably, we found more than 65% of β-Gal neurons were also Fos-positive, comparable to prior studies (Bossert et al., 2012, Fanous et al., 2012; Funk et al., 2016).

## 3. Results

### 3.1. Histology

Location of injector tips were within or directly above the BLA as shown in Fig 1. Final group numbers based on proper cannula placements were CS+Daun02 (n=12 total; n=6 for Reversal Session 1 [R1]; n=6 for Reversal Session 15 [R15]), CS−Daun02 (n=11 total; n=5 for R1; n=6 for R15), and Vehicle (n=19 total; n=10 for R1; n=9 for R15).

**Figure 1.**
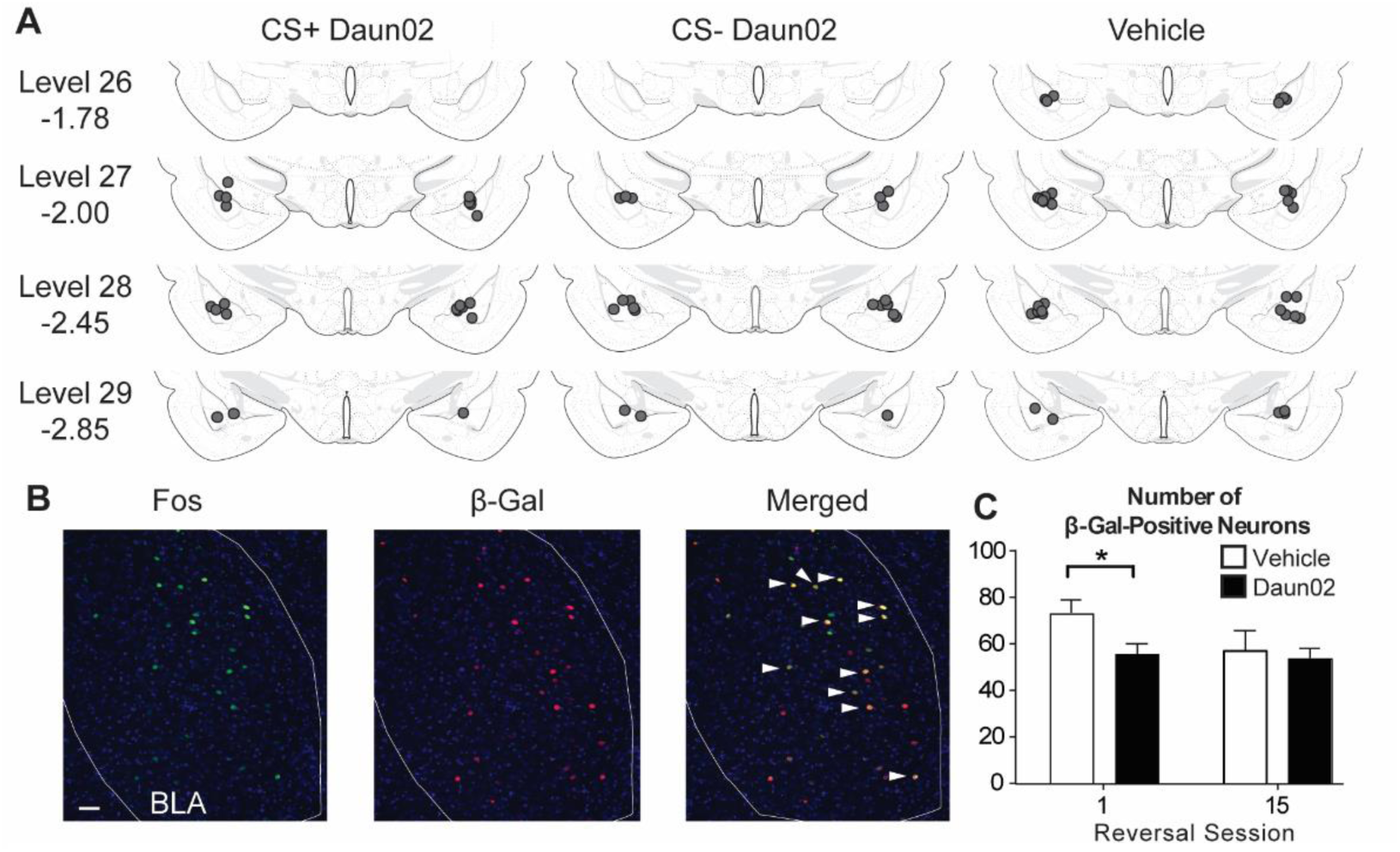
Cannula placements and Fos and β-Gal induction. (**A**) All cannula placements were within levels 26-29 of the BLA (−1.78 to −2.85mm from Bregma as indicated) and similar across drug groups. (**B**) Representative images showing Fos, β-Gal, and colocalization (white arrows). (**C**) There was a significant reduction in β-Gal-labeled neurons in the group that received Daun02 and were perfused after the first reversal session, but not the last reversal session. Scale bar = 25 μm. * *p* < 0.05.

To verify Daun02 inactivation methodology, the number of β-Gal-labeled neurons was compared between drug treatment groups. Our *a priori* hypothesis was a decrease in the number of β-Gal-labeled neurons in rats that received the Daun02 compared to Vehicle, specifically in the groups that received only 1 session of reversal learning (Fig. 1B, C), and not 15 sessions. This was statistically confirmed (R1: *t*(19) = −2.459, *p*=0.02; R15: *t*(19) = −0.442, *p*>0.5). No difference was found between the R15 groups, suggesting additional ensembles were activated as new learning occurred to potentially compensate for the initial neuronal inactivation.

### 3.2. Discriminative Conditioning and Induction Session

All groups successfully discriminated between the CS+ and CS−, as shown by higher conditioned responding and faster latencies to the CS+ compared to the CS− during the tenth training session (data not shown). These results were expected since no drug was given during training, and group allocation was based on drug treatment after the induction session. A group (CS+Daun02, CS−Daun02, Vehicle) X CS (CS+, CS−) repeated measures ANOVAs showed an effect of CS Elevation on food cup behavior (*F*(1,39) = 331.00, *p* < 0.001) and an effect of CS on latency (*F*(1,39) = 110.85, *p* < 0.001), but no Group effects or interactions (*F*’s < 0.5). After discriminative conditioning, rats underwent an induction session with presentation of either the CS+ or CS− (not both) to induce Fos in the BLA in response to the respective CS. Conditioned responding was similar to the last conditioning session with higher responding in the CS+ induction groups compared to the CS− induction groups (data not shown). A Drug (Daun02, Vehicle) X CS Elevation (CS+, CS−) ANOVA during the induction session confirmed an effect of CS (*F*(1,38) = 61.50, *p* < 0.001), but no effect of drug or interaction (*F*’s < 2.3, *p*’s > 0.1). These results were expected since drug infusions occurred after the induction session and confirm similar responding between drug groups within their respective CS induction.

### 3.3. Reversal Learning

#### 3.3.1. Reversal Session 1

The group that received Daun02 following CS+ presentations during the induction session (CS+Daun02), and thus CS+ neuronal ensemble inactivation, had lower conditioned responding to the same CS, now rCS−, during reversal session 1 (Fig. 2A). Analysis of conditioned responding with a Group X CS Elevation repeated measures ANOVA showed a significant effect of CS (*F*(1,39) = 273.57, *p* < 0.001), but no effect of Group (*F* = 1.80, *p* = 0.18) or interaction (*F* = 2.19, *p* = 0.13). Simple effect analyses on each rCS showed a Group effect on average responding to the rCS− (*F*(2,39) = 4.08, *p* = 0.025), with lower responding in the CS+Daun02 group compared to the CS−Daun02 (*p* < 0.01) and Vehicle (*p* = 0.05) groups, and no difference between the CS−Daun02 and Vehicle groups (*p* > 0.1). No group differences were found in rCS+ responding (*F* < 0.4, *p* > 0.5).

**Figure 2.**
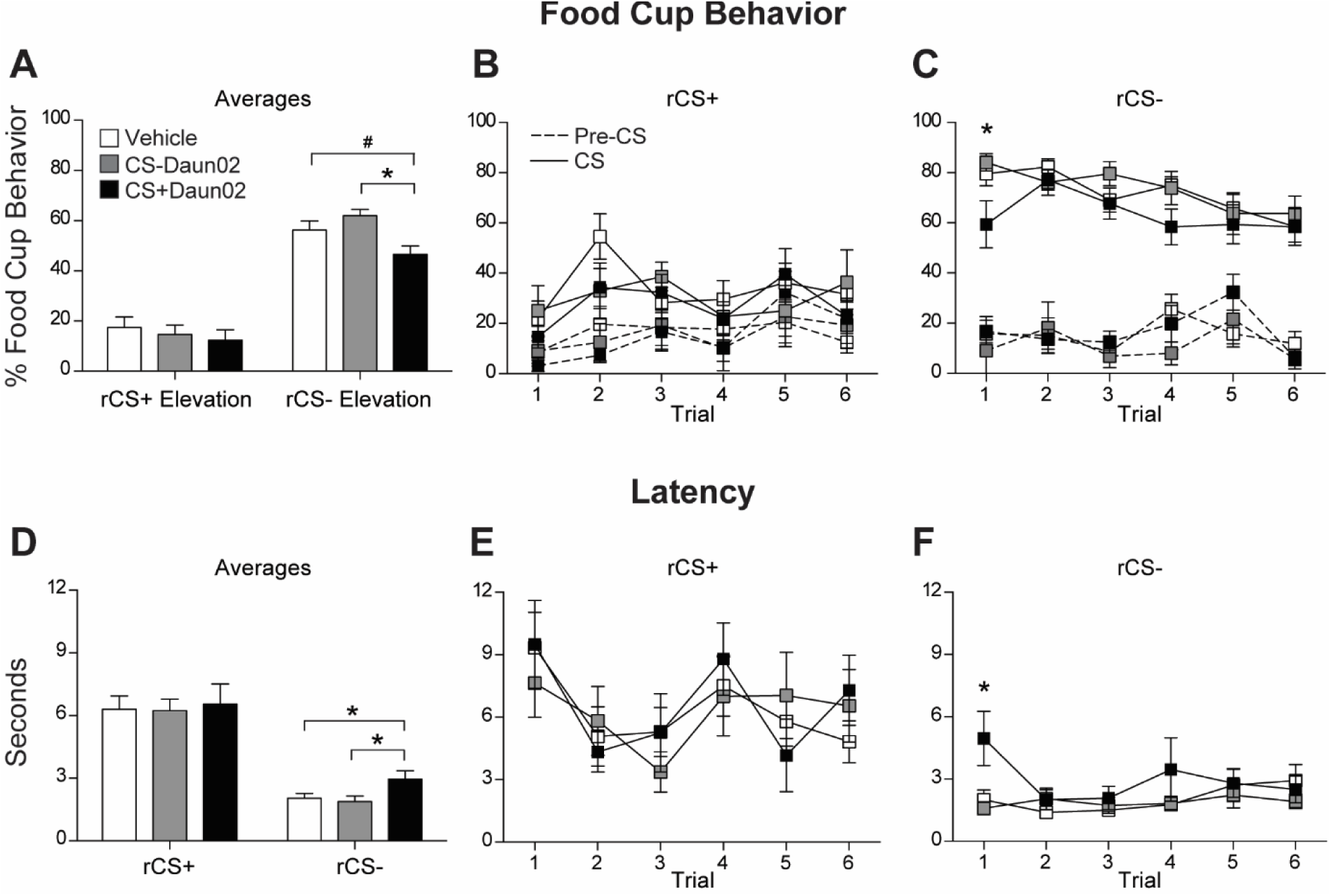
Conditioned responding during Reversal Session 1. (**A**) Average food cup responding (mean ± SEM) during rCS+ and rCS− during Reversal Session 1. Data shown as Elevation score (CS responding minus pre-CS [baseline] responding). (**B**,**C**) Food cup responding to each rCS+ trial (**B**) and rCS− trial (**C**) during the session. Solid line represents responding during the cue, and dashed line represents responding during the pre-CS (baseline) period. (**D**) Average latency to approach the food cup (mean ± SEM) during the rCS+ and rCS− during the session. (**E**,**F**) Latency responding to each rCS+ trial (**E**) and rCS− trial (**F**) during the session. # *p* = 0.05; * *p* < 0.05.

To evaluate the recall of the cue value following Daun02 neuronal ensemble inactivation, we analyzed each trial in the first reversal session, with emphasis on the first trial. The CS+Daun02 group showed lower conditioned responding the first time the CS+, now rCS−, was presented (Fig. 2C). There was a main effect of Group (*F*(1,39) = 3.93, *p* = 0.028) with the CS+Daun02 group showing lower conditioned responding to the CS−Daun02 (*p* = 0.015) and Vehicle (*p* = 0.023) groups.

Additionally, we analyzed latency to approach the food cup after the onset of each cue and found rats approached the food cup faster during the rCS− compared to the rCS+, as expected since the rCS− was previously the CS+. A Group X CS ANOVA showed an effect of CS on overall average latency to respond to the food cup (*F*(1,39) = 125.36, *p* < 0.001; Fig. 2D) indicating rats approached the food cup faster during the rCS− compared to the rCS+, but there was no Group effect or interaction (*F*’s < 1, *p*’s < 0.5). Simple effect analyses on each rCS showed a Group effect on latency to the rCS− (*F*(2,39) = 3.61, *p* = 0.037), with slower latencies in the CS+Daun02 group compared to the CS−Daun02 (*p* = 0.021) and Vehicle (*p* = 0.026) groups, and no difference between the CS−Daun02 and Vehicle groups (*p* > 0.1). No group differences were found in latency to the rCS+ (*F* < 0.1, *p* < 0.5).

Again, we analyzed the first trial during the first reversal session for latency to the rCS− and found a main effect of Group (*F*(2,39) = 5.27, *p* < 0.01) with the CS+Daun02 group showing slower latencies to the rCS− compared to the CS−Daun02 and Vehicle groups (*p*’s < 0.01; Fig. 2F). We found no differences for other rCS− trials or rCS+ trials (*p*s>0.1).

#### 3.3.2. Reversal Learning Across Sessions

We analyzed responding in a group of rats that underwent 15 sessions of reversal learning to determine if inactivation of specific CS BLA neuronal ensembles interfered with updating the new values of the cues during reversal learning (Fig. 3). The group that received Daun02 following CS+ induction showed a decrease in conditioned responding to the same CS, now the rCS−, throughout reversal learning (Fig. 3B). Analysis of responding to CSs with a Group X CS Elevation X Session repeated measures ANOVA showed CS Elevation X Group interaction (*F*(2,18) = 7.33, *p* < 0.01), and expected main effects of Session (*F*(3,54) = 5.23, *p* < 0.01) and CS (*F*(1,18) = 40.82, *p <* 0.001) and a Session X CS Elevation interaction (*F*(3,54) = 81.38, *p* < 0.001), but no other effects or interactions (*F*’s < 2, *p* > 0.1). Follow-up analyses confirmed a significant effect of Group on conditioned responding to the rCS− (*F*(2,18) = 5.08, *p* = 0.018) with the CS+Daun02 group showing lower conditioned responding to the rCS−, which was previously their CS+, across reversal learning compared to the CS−Daun02 (*p* = 0.010) and Vehicle (*p* = 0.014) groups. The decrease was specifically during reversal session 1 as described above and session 15 (*F*(2,18) = 4.71, *p* = 0.023), where the CS+Daun02 group had significantly lower responding compared to the Vehicle and CS−Daun02 groups (*p*’s < 0.015).

**Figure 3.**
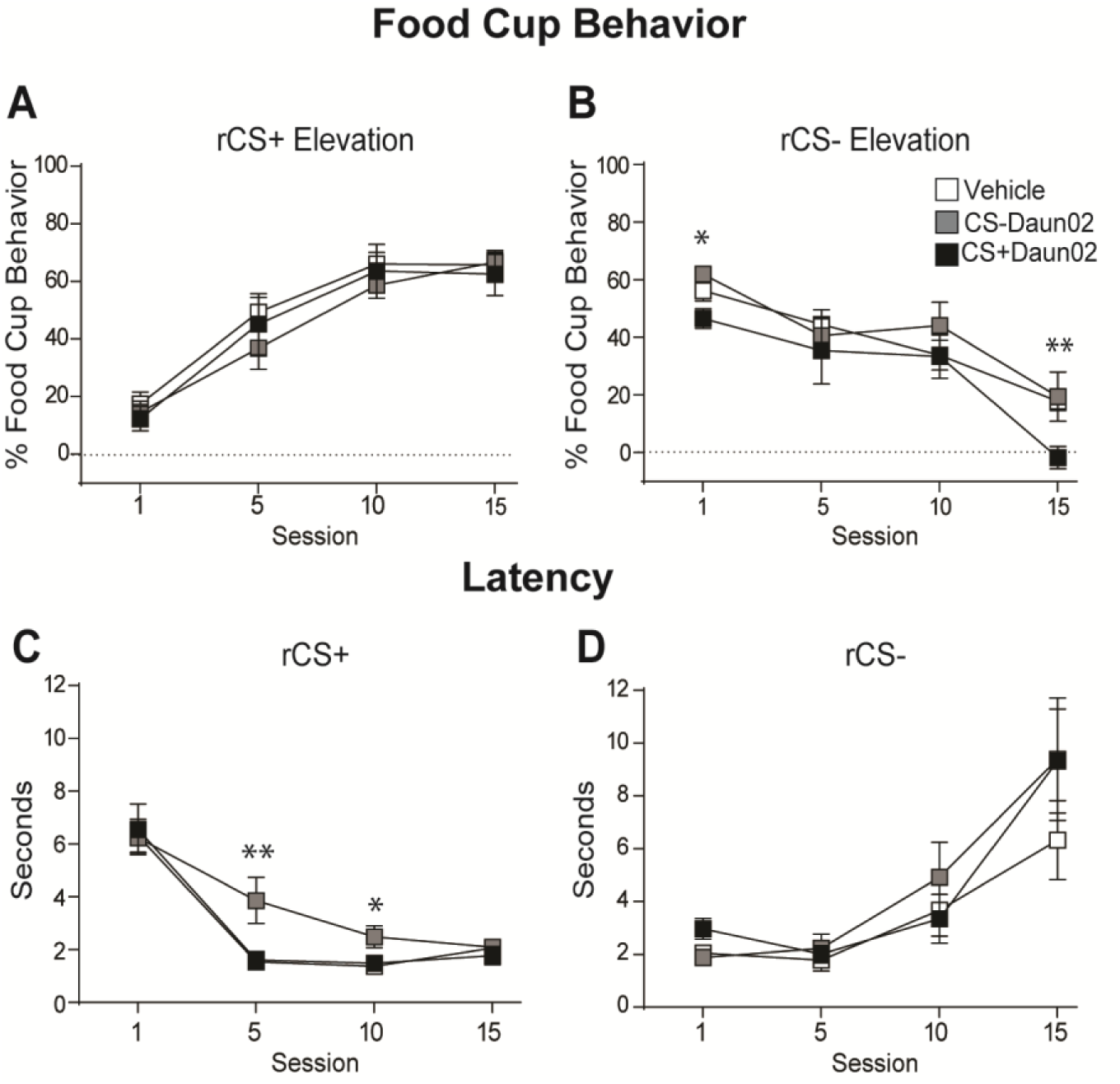
Conditioned responding throughout reversal learning. (**A**,**B**) Average food cup responding (mean ± SEM) during rCS+ (**A**) and rCS− (**B**) throughout reversal learning. Data shown as Elevation score (CS responding minus pre-CS [baseline] responding). (**C**,**D**) Average latency to approach the food cup (mean ± SEM) during the rCS+ (**C**) and rCS− (**D**) throughout reversal learning. * *p* < 0.05; ** *p* < 0.015.

#### 3.3.3.

The analyses on latency responding showed the group that received Daun02 following CS− induction was slower to respond to the food cup after the same cue presentation, now rCS+, during reversal learning (Fig 3C). A Group X CS X Session repeated measures ANOVA found expected main effects of Session (*F*(3,54) = 14.51, *p* < 0.001) and CS (*F*(1,18) = 6.85, *p* < 0.02), and a Session X CS interaction (*F*(3,54) = 40.79, *p* < 0.001), but no Group effects or interaction (*F*’s < 1.5, *p*’s > 0.1). Simple effect analyses showed an effect of Group on responding to the rCS+ during session 5 (*F*(2,18) = 7.39, *p* < 0.01) and session 10 (*F*(2,18) = 4.21, *p* < 0.05). The CS−Daun02 group had significantly longer latencies to the rCS+ during session 5 compared to the CS+Daun02 and Vehicle groups (*p*’s < 0.01) and session 10 compared to the CS+Daun02 group (*p* < 0.01) and Vehicle group (*p* = 0.037). We found no differences in latency to respond to the rCS− (*p*’s>0.1).

## 4. Discussion

The current study examined if potentially separate BLA neuronal ensembles that are activated during CS+ and CS− memory recall are necessary for value updating when the CS contingencies are reversed. This was accomplished by testing if inactivating the neuronal ensemble that responds to a particular learned CS altered the memory of that specific CS and not the other CS. Additionally, we examined if CS-specific neuronal ensembles are necessary to learn the new associations to the same CS when the outcome is changed during reversal learning. Using the Daun02 method, we specifically inactivated BLA neuronal ensembles that were activated by either a CS that was previously associated with food (CS+) or a CS not paired with food (CS−), and then evaluated conditioned responding when the outcomes of the cues were switched during reversal learning. We found that the group that received the Daun02 following the CS+ session (CS+Daun02 group) showed a decrease in conditioned responding to the same CS, now the rCS−, during the first reversal session and specifically during the first presentation of that CS prior to any new information about the outcome. This decreased conditioned responding to the rCS− indicates that the CS+ neuronal ensemble was necessary to recall the previously learned outcome of the cue during the first reversal session and to incorporate this information during reversal learning. Second, we found that the group that received Daun02 following CS− induction (CS−Daun02 group) was slower to approach the food cup (i.e. longer latency) after the same cue presentation, now the rCS+, during reversal learning, specifically during reversal sessions 5 and 10. Together, these results support our hypotheses that separate BLA neuronal ensembles mediate CS+ and CS− memory recall, and reactivation of each cue-specific neuronal ensemble is necessary for updating the value of that specific learned cue, in order to respond appropriately during reversal learning. Importantly, these results also indicate plasticity of BLA neuronal ensembles when cues’ values are altered since all rats eventually showed the same levels of conditioned responding by the end of reversal learning.

The observed impairments in conditioned responding were specific to the CS to which the neuronal ensemble was inactivated and did not cause general impairments in behavioral responding. The CS+Daun02 group was impaired on responding to rCS−, previously the CS+, and CS−Daun02 group was impaired on responding to rCS+, previously the CS−. This suggests that our preparation inactivated separate CS+ and CS− neuronal ensembles, which impaired subsequent, CS-specific reversal learning. This is in agreement with prior work that found specific effects of neuronal ensembles inactivation by Daun02 (Pfarr et al., 2015). Additionally, other studies have shown altered reward-seeking behaviors due to specific neuronal ensemble inactivation with this method (Caprioli et al., 2017; Cole et al., 2020; Cruz et al., 2014; de Guglielmo et al., 2016; Fanous et al., 2012; Funk et al., 2016; George and Hope, 2017; Koya et al., 2009; Pfarr et al., 2015; Warren et al., 2016; Whitaker et al., 2017; Xue et al., 2017; (Josselyn and Frankland, 2018; Josselyn and Tonegawa, 2020).

The results of the current study are in agreement with previous studies that showed separate BLA neuronal ensembles respond to distinct learned cues. Previous studies have shown that ∼60% of BLA neurons respond to a distinct learned cue during appetitive learning, and then half of these neurons alter their responding when the outcomes are switched during reversal learning in rats (Schoenbaum et al., 1999; Zhang and Li, 2018) and primates (Paton et al., 2006). Interestingly, in both studies the neurons responded selectively to specific cues prior to correct behavioral performance, indicating BLA neurons are tracking the outcome and the value of learned cues to guide behavior. Similarly, another study showed a subset of BLA neurons (∼10%) respond to a well-learned reward predictive cue, but then distinctly alter their responses when food is no longer delivered during extinction (“reinforcement-omission” neurons; (Tye et al., 2010)), suggesting this subset of neurons may be tracking the outcome of the learned cues. This indicates that BLA neurons can alter their responses based on environmental changes, signifying neural plasticity.

The current results indicate that the BLA regulates the updating of the value of learned appetitive cues based on the current outcome. This was confirmed for both groups that received cue-specific neuronal ensemble inactivation by Daun02: the CS+Daun02 group had lower conditioned responding to the same CS during reversal learning (rCS−), and the CS−Daun02 group was slower to approach the food cup after presentation of the same CS during reversal learning (rCS+). Indeed, previous studies have shown that an intact BLA is needed to access the value of the learned cue in order to appropriately update it when the outcome is changed and alter behavioral responding (as reviewed in (Wassum and Izquierdo, 2015)). The BLA encodes the value of the cues during learning (Cole et al., 2013; Esber and Holland, 2014; Parkes and Balleine, 2013; Piette et al., 2012; Schoenbaum et al., 1999; Tye and Janak, 2007; Uwano et al., 1995) and is involved in appetitive cue discrimination (Ambroggi et al., 2008; Ishikawa et al., 2008) and reversal learning (Churchwell et al., 2009). However, several studies have shown the BLA may not be critical for initial acquisition of cue value learning (Balleine et al., 2003; Corbit and Balleine, 2005; Hatfield et al., 1996; Holland et al., 2002; Parkinson et al., 2000), but it is critical to encode and assess the representation of the learned associations to alter subsequent behavioral motivation and learning (Blundell et al., 2001; Corbit and Balleine, 2005; Coutureau et al., 2009; Everitt et al., 2003; Galarce et al., 2010; Hatfield et al., 1996; Holland et al., 2002; Holland and Petrovich, 2005; Johnson et al., 2009; Ostlund and Balleine, 2008; Petrovich, 2013; Setlow et al., 2002; Tye and Janak, 2007; Wassum and Izquierdo, 2015; Hoang and Sharpe, 2021; Fisher et al., 2020). This suggests a specific role for the BLA in reward value representation when appetitive learning is altered, in agreement with the current findings.

## 5. Conclusions

The current study investigated the plasticity across a learning paradigm that requires value updating. We found that inactivation of the BLA neuronal ensemble responsive during a specific learned cue recall resulted in impaired conditioned responding to the same cue during reversal learning without interfering with responding to the other learned cue. These results show distinct neuronal ensembles within the BLA are activated during specific cue memory recall and are necessary to update the value of that cue during reversal learning.

## Acknowledgements

We thank Bret Judson and the Boston College Imaging Core for infrastructure and support, and Dr. Bruce Hope and Dr. Francois Vautier for advice and protocols regarding breeding *c-fos-lacZ* transgenic rats. We also thank the Boston College undergraduate researchers who helped with numerous aspects of this project.

## Supplementary Materials and Methods

### Subjects

Experimentally naïve male and female *Fos-lacZ* transgenic rats bred in the animal facility at Boston College were used. Rats had *ad libitum* access to food (standard laboratory chow) and water and were grouped housed until ∼2 days prior to surgical procedures when they were individually housed and acclimated to experimenter handling (∼2 months old). The colony room was maintained on a 12-hour light/dark cycle with lights on at 06:00. All procedures complied with the National Institutes of Health *Guidelines for Care and Use of Laboratory Animals* and were approved by the Boston College Animal Care and Use Committee.

### Surgical Procedure

For surgeries, rats were anesthetized with either isoflurane (1-3%) or a mixture (1 ml/kg body weight) of ketamine (50 mg/ml) and xylazine (10 mg/ml). Rats were bilaterally implanted with 22- or 26-gauge guide cannulas (Plastics One Inc.) targeting the BLA. The flat-skull coordinates from bregma were 2.7-3.0 mm anteroposterior, ±4.6-4.8 mm mediolateral, and −7.1 mm dorsoventral. Cannulas were anchored to the skull with screws and dental cement. Obturators were inserted into the guide cannulas where they remained throughout the experiment, except during microinfusions. Triple antibiotic cream was applied around the cannula cap, and rats were given analgesic Rimadyl (Henry Schein; 50 mg/ml) in a sterile saline solution (4.4 mg/kg) the day of surgery and chewable Rimadyl tablets (Bio Serv; 2 mg/1 tablet/100 g bodyweight) for two days after surgery. Rats were allowed to recover for at least 5 days post-surgery prior to behavioral training and were monitored and weighed daily.

### Intracranial Infusions

For microinfusions, obturators were removed, and injectors were inserted into the guide cannula. Injectors that projected 1 mm ventral to the tip of the guide cannula were connected via polyethylene tubing to 10 μl Hamilton syringes and mounted onto an infusion pump. Either Daun02 or vehicle solution were infused at a rate of 0.5 μl/side over 1 min with an additional 1 min post-injection diffusion time. Following infusions, obturators were reinserted, and rats were returned to their home cage.

Daun02 (Sequoia Research Products) was dissolved in 5% dimethyl sulfoxide (Sigma-Aldrich), 6% Tween-80, and 89% phosphate buffered saline. Vehicle consisted of the same solution without Daun02.

### Apparatus

All habituation and training occurred within the same set of behavioral chambers (30 × 28 × 30 cm; Coulbourn Instruments) located in a separate room from the colony room. The top and sides of the chambers were aluminum, and one side contained a recessed food cup (3.2 × 4.2 cm). The front and back of the chamber were transparent Plexiglas, and the front panel was hinged. The floor was composed of 5 mm stainless-steel rods spaced 15 mm apart. Each chamber was contained within an isolation cubicle (79 × 53 × 53 cm; Coulbourn Instruments) composed of monolithic rigid foam walls and a ventilation fan (55 dB). On the rear wall of each isolation cubicle was a video camera connected to a recording system (Coulbourn Instruments).

The conditioned stimuli (CSs) were a 10s 75dB, 2kHz tone and a 10s 75dB white noise. The unconditioned stimulus (US) was two food pellets (TD pellets; formula 5TUL, 45 mg: TestDiet) that were delivered into the food cup of each chamber when applicable. GraphicState 3.0 software system (Coulbourn Instruments) controlled all stimuli.

### Behavioral Training Procedure

Rats were food restricted to maintain 90% of their maximum recorded body weight throughout training. Two days prior to training, rats received 1g of the US in their home cage to reduce novelty to the training food. The following day rats underwent a ∼30 min magazine training in the behavioral chambers where they were given random deliveries of the US to familiarize them with eating from the food cup.

#### Discriminative Conditioning

Rats received ten 30 min training sessions over 5 days (2 sessions/day). Each session consisted of intermixed presentations of two different auditory cues, each presented six times. One auditory CS (e.g. tone) was immediately followed by delivery of 2 TD pellets (CS+), and the other CS (e.g. noise) was presented alone (CS−). Auditory cues that served as CS+ and CS− were counterbalanced. The inter-trial intervals (ITIs) were between 60-219s, and ITIs and CS order varied randomly across training sessions.

#### Induction session

Following successful discriminative conditioning, rats underwent an induction session in which they were given six presentations of either the CS+ or CS− (without US) to induce expression of the *c-fos* gene (Fos protein) in neuronal ensembles activated during that session. The transgene *lacZ* produces the protein β-galactosidase (β-gal), and its expression is controlled by *c-fos* gene. Thus, neurons that produce *c-fos* in response to a stimulus also produce β-gal (Koya et al., 2009; Cruz et al., 2013). Ninety minutes following the beginning of the induction session when Fos expression is optimal, rats were briefly anesthetized with isoflurane and received infusions of either Daun02 or vehicle. Infusions of Daun02, which is catalyzed by β-gal into daunorubicin, results in a reduction in neuronal excitability (Santone et al., 1986; Engeln et al., 2016) and eventually in cell death by apoptosis (Pfarr et al., 2015). Therefore, with this method, we targeted specific neurons that were activated (and expressed Fos) during the cue presentations (CS+ or CS−).

#### Reversal Learning

At least three days following induction session (optimal time for Daun02 inactivation [Koya et al., 2009]), rats began reversal learning. These sessions were similar in length and number of CS presentations as the discriminative conditioning sessions; however, the outcomes of the CS+ and CS− were reversed. The original cue that was associated with the TD pellets (CS+) was now no longer followed by the pellets (reversal CS−; rCS−), and the cue that was previously followed by nothing (CS−) was now always followed by the delivery of pellets (reversal CS+; rCS+). A subset of rats underwent only 1 reversal session to observe responding to the initial (“true”) memory recall of the CSs and for induction of Fos and β-gal, and the rest of the rats underwent 15 sessions of reversal learning (1-2 sessions/day).

### Behavioral Measures

The primary measure of learning was conditioned responding to the food cup during the presentations of the CSs. This behavior was defined as rats standing in front of and directly facing the food cup or demonstrating distinct nose pokes into the food cup. Trained observers unaware of group allocation recorded rats’ behavior every 1.25 seconds during each 10 second CS, as well as during 10 seconds prior to the onset of the CS as a measure of baseline responding (pre-CS). Only one behavior was recorded at each observation (food cup or nothing). The total number of identified food cup observations for each CS during each period was summed and converted into a percentage of total time rats spent in the food cup, which was used to calculate the mean value for each group. CS elevation was calculated by subtracting pre-CS responding from CS responding (CS minus pre-CS) and then the mean value was calculated for each group.

Latency was measured as the time elapsed from the onset of the CS until the rat approached the food cup within 20s from the CS onset (10s CS plus 10s to collect the US, if applicable). After this time, behavior was considered unspecific to the presentation of the CS, and a maximum latency of 20s was assigned to any trial that surpassed this time without a response. For each rat, latency for each trial was used to calculate the average latency responding for each CS during each session and then used to calculate the mean value for each group.

### Histological Procedures and Immunohistochemistry

Ninety minutes following either the first or 15^th^ reversal session, rats were perfused, and brain tissue was collected for analysis of Fos and β-gal induction and cannula placement verification. Rats were given a lethal dose of tribromoethanol (375 mg/kg; 1.5ml/100g bodyweight; i.p.) and transcardially perfused with 0.9% saline followed by ice cold 4% paraformaldehyde in 0.1M borate buffer. Brains were stored overnight (18-24hr) in the paraformaldehyde solution with 12% sucrose at 4°C, and then rapidly frozen in hexanes cooled in dry ice and stored at −80°C.

Frozen brains were sliced into 30 μm coronal slices using a sliding microtome (Leica Biosystems) and collected into four serially adjacent series. One series of tissue was mounted from a potassium phosphate-buffered saline solution (KPBS) onto gelatin-coated slides, dried at 45°C, dehydrated through graded alcohols, stained with thionin, cleared in xylenes, and coverslipped with DPX Mountant. The thionin stain allowed for identification of neuroanatomical borders and cannula placements, which were examined under a light microscope and mapped using a rat brain atlas (Swanson, 2004).

Another series of tissue was stained for identification of Fos and β-gal induction to verify the Daun02 inactivation method (i.e. decreased number of β-gal positive neurons in Daun02 infused groups compared to vehicle infused groups). Brain tissue was incubated with anti-Fos primary antibody raised in rabbit (1:10,000, ABE457; Millipore, Temecula, CA, USA) and anti-β-gal primary antibody raised in mouse (1:2,000, sc-65670; Santa Cruz, Dallas, TX, USA) in a solution containing KPBS, 2% normal donkey serum (NDS; Jackson ImmunoResearch, West Grove, PA, USA), and 0.3% Triton X-100 (Sigma-Aldrich) for 72 hours at 4°C. Tissue was then rinsed with KPBS and incubated with secondary fluorescent antibodies: Alexa 488 anti-rabbit raised in donkey (1:200, A21206; Invitrogen, Carlsbad, CA, USA) and Alexa 594 anti-mouse raised in donkey (1:200, A21203, Invitrogen) in KPBS, NDS, and Triton X-100 for 1 hr in semi-darkness. Tissue was rinsed again and mounted onto Superfrost slides in semi-darkness, air-dried, coverslipped with Vectashield Hardset Mounting Medium with DAPI (4’,6-diamidino-2-phenylindole; H-1500; Vector Laboratories), and stored at 4°C until analysis.

### Image Acquisition and Analysis

Images throughout the BLA were acquired with a Zeiss Axio Image Z2 fluorescence microscope (Carl Zeiss Microscopy GmbH, Jena, Germany) and attached Hamamatsu ORCA-R2 camera (Bridgewater, NJ, USA) using Zen software. Images were pseudo-colored with green for Fos, red for β-gal, and blue for DAPI (nuclear counterstain). Single Fos-labeled and single β-gal-labeled neurons were manually counted from acquired images and summed for each stain for each rat to calculate the total number of Fos-labeled neurons and the total number of β-gal-labeled neurons. Each count was then averaged within groups to determine mean counts for each group.

### Statistical Analysis

Behavioral data were analyzed using a mixed design analysis of variance (ANOVA) with Group (CS+Daun02, CS−Daun02, Vehicle) as a between-subjects factor and type of CS Elevation (CS+, CS− or rCS+, rCS−) and conditioning sessions (discriminative conditioning session 10 or reversal learning session 1, 5, 10, and 15) as within-subjects repeated factors. The dependent variable was the percentage of time rats displayed conditioned responding and latency (in seconds) for all behavioral training. For the neural data analysis, the dependent variable was the number of β-gal-labeled neurons across experimental groups. A significance value of *p* < 0.05 was used for all analyses. SPSS software was used for all statistical analyses.

Daun02 groups were treated as separate groups due to the nature of the hypothesis—the CS+ and CS− are mediated by separate BLA neuronal ensembles and inactivating the neuronal ensembles that respond to a particular CS will only alter responding of that specific CS and not the other CS. Furthermore, in order to confirm Daun02 did not have a general drug effect regardless of CS induction, statistical analyses compared all Daun02 infused rats to Vehicle infused rats and found no overall differences caused specifically by the drug for any measures recorded (*p*s > 0.05). This analysis shows that any observed behavioral differences in either Daun02 group is not due to a general effect of the drug but is instead due to the specific CS neuronal ensemble inactivation caused by Daun02 (see Pfarr et al., 2015).

Additionally, groups that received Vehicle infusions following either CS+ or CS− induction session were combined into one group for their respective reversal session analyses. The rationale was that responding should be similar between these groups since no neural deficits occurred due to infusing the vehicle solution.

